# Self-fertilization, but not mating strategy, predicts the evolution of sex allocation plasticity in a hermaphroditic flatworm genus

**DOI:** 10.1101/2020.06.12.149351

**Authors:** Pragya Singh, Lukas Schärer

**Author notes:** Chemical Ecology, Bielefeld University, Universitätsstr. 25, 33615 Bielefeld, Germany. **Author Contributions:** PS and LS designed the study, analysed the data and wrote the manuscript. PS performed the experiment. All authors read and approved the final manuscript. **Ethical note:** All animal experimentation was carried out in accordance to Swiss legal and ethical standards. **Data accessibility:** All data in this manuscript will be deposited online (xx).

## Abstract

Sex allocation (SA) theory in simultaneous hermaphrodites predicts that optimal SA is influenced by local sperm competition (LSC), which occurs when related sperm compete to fertilize a given set of eggs. Different factors, including the mating strategy and the ability to self-fertilize, are predicted to affect LSC and hence the optimal SA. Moreover, since the LSC experienced by an individual can vary temporally and spatially, this can favour the evolution of SA plasticity. Here, using seven species of the free-living flatworm genus *Macrostomum*, we document sizable interspecific variation in SA, but neither their mating strategy nor their ability to self-fertilize significantly predicted SA among these species. Since we also found considerable interspecific variation in SA plasticity, we further estimated standardized effect sizes for plasticity in response to i) the presence of mating partners (i.e. in isolation vs. with partners) and ii) the strength of LSC (i.e. in small vs. large groups). We found that self-fertilization predicted SA plasticity with respect to the presence of mating partners, with plasticity being lower for self-fertilizing species. Finally, we showed that interspecific variation in SA is higher than intraspecific variation due to SA plasticity. Our study suggests that both SA and SA plasticity are evolutionarily labile, with self-fertilization predicting the latter in *Macrostomum*.

## Introduction

Sex allocation (SA) theory in simultaneous hermaphrodites predicts the optimal allocation of resources towards the male versus female function (Charnov 1982). Local sperm competition (LSC), which can be thought of as the inverse of sperm competition (Parker 1970, 1998), occurs when related sperm (usually from the same individual) compete for access to a given set of eggs, and it can favour a female-biased optimal SA in simultaneous hermaphrodites (Schärer 2009; Schärer and Pen 2013). LSC can be considered analogous to local mate competition (Hamilton 1967), i.e. the competition between related males for access to mates, which can lead to a female-biased sex ratio in species with separate sexes. However, LSC does not require mates to be related or a spatial population structure to be present, since the local competition occurs among competing sperm, not competing males. An example of a factor that can affect LSC, and hence the optimal SA, is the mating group size (Charnov 1980, 1982), denoted as *K*+1, where *K* is the number of sperm donors from which a sperm recipient receives sperm at the time its eggs are fertilized. Specifically, Charnov’s mating group size model predicts that the optimal proportion of resources an individual allocates to the male function should increase with the number of sperm donors that are present in a local mating group.

The link between optimal SA and LSC can be visualised in terms of male fitness gain curves, which describe how much fitness is gained through the male function per unit resource investment (Charnov 1982; Lloyd 1984; Schärer 2009). Under monogamy, LSC is necessarily very high, since the competing sperm are maximally related to each other, resulting in the male fitness gain curve saturating quickly. Thus, any investment into the production of more sperm than are required to fertilize the partner’s eggs is a waste of resources, as the related sperm simply compete amongst themselves for fertilizations. Instead, these resources could be more profitably invested into the individual’s own female function (Charnov 1982), since the female fitness gain curve is often assumed to be linear (Rademaker and de Jong 1999; Campbell 2000; Schärer 2009), and may sometimes even be accelerating (Rosas and Domínguez 2009). Consequently, under monogamy, a simultaneous hermaphrodite is expected to have a highly female-biased SA (Charnov 1980, 1982). But as the mating group size increases—i.e. with more competitors contributing sperm to a given recipient—an individual’s sperm are more and more competing with unrelated sperm. So, it now pays off to invest into the male function to gain a greater share of the fertilizations, and the male fitness gain curve is predicted to linearize, with a subsequent shift towards a more equal SA (Charnov 1982; Schärer 2009; Schärer and Pen 2013).

An interesting question that arises then is whether and how interspecific variation in reproductive biology, including different mating strategies or the ability to self-fertilize, could influence the strength of LSC and hence the optimal SA. For example, hermaphrodites often evolve different mating strategies in response to sexual conflict over mating roles, i.e. over whether to mate as a sperm donor and/or as a sperm recipient (Charnov 1979; Michiels 1998; Schärer et al. 2015). One such mating strategy is reciprocal mating (also called reciprocal copulation), in which both partners mate in both mating roles, and thus donate and receive sperm simultaneously. This strategy can lead to individuals being quite willing to engage in mating in order to get an opportunity to donate sperm, and this general willingness likely results in reduced precopulatory sexual selection. The ensuing higher mating rates could in turn result in increased sperm competition (and thus a decrease in LSC) and more intense postcopulatory sexual selection. Interestingly, however, the presence of certain postcopulatory processes could also increase LSC (Schärer 2009). Indeed, theoretical studies have predicted that different postcopulatory sexual selection processes, such as sperm displacement, sperm digestion, and cryptic female choice, can lead to the removal of sperm of one or multiple donors from competition, resulting in changes in LSC and hence the predicted optimal SA (Charnov 1996; Greeff and Michiels 1999; Pen and Weissing 1999; van Velzen et al. 2009; Schärer and Pen 2013).

In addition, it has been suggested that postcopulatory sexual selection and sexual conflict among simultaneous hermaphrodites can also favour the evolution of mating strategies that allow donors to bypass these postcopulatory processes and fertilize the eggs more directly (Charnov 1979; Michiels 1998). One such mating strategy is forced unilateral insemination, in which one partner mates only in the male role and donates sperm, e.g. via traumatic or hypodermic insemination (Charnov 1979; Lange et al. 2013; Reinhardt et al. 2015), while the other partner only mates in the female role, likely often against its interests. It is currently difficult to predict the direction and magnitude of changes in LSC resulting from such a shift in mating strategy—and the effects that such shifts may thus have on optimal SA—since we still know relatively little about the particular postcopulatory processes involved in these mating strategies. In species with reciprocal mating there may be some control over who or how many partners an individual receives and stores sperm from, and there could be strategies that lead to the (conceivably selective) removal of received sperm (van Velzen et al. 2009), possibly leading to increased LSC. In species with hypodermic insemination, however, such control might be more limited, since hypodermic sperm might function in a less easily controlled environment, with sperm from multiple donors thus mixing more randomly in a fair-raffle-type sperm competition. This would be expected to lower the strength of LSC (Schärer and Janicke 2009).

In addition to the contrast between reciprocal copulation and hypodermic insemination, species that employ self-fertilization might also differ in optimal SA from species that primarily outcross, because self-fertilization greatly increases the strength of LSC. Thus, self-fertilization leads to diminishing fitness returns from investing in the male function, favouring the evolution of a more female-biased optimal SA compared to outcrossing species (Charlesworth and Charlesworth 1981; Charnov 1982, 1987), which is supported by empirical work in both plants and animals (Lemen 1980; Schoen 1982; McKone 1987; Johnston et al. 1998; Lemen 1980). In general, we would expect any process that has an effect on LSC to affect the shape of the male fitness gain curve. And this in turn leads to different predictions for the optimal SA (Schärer 2009), as shown in Figure 1.

**Figure 1.**
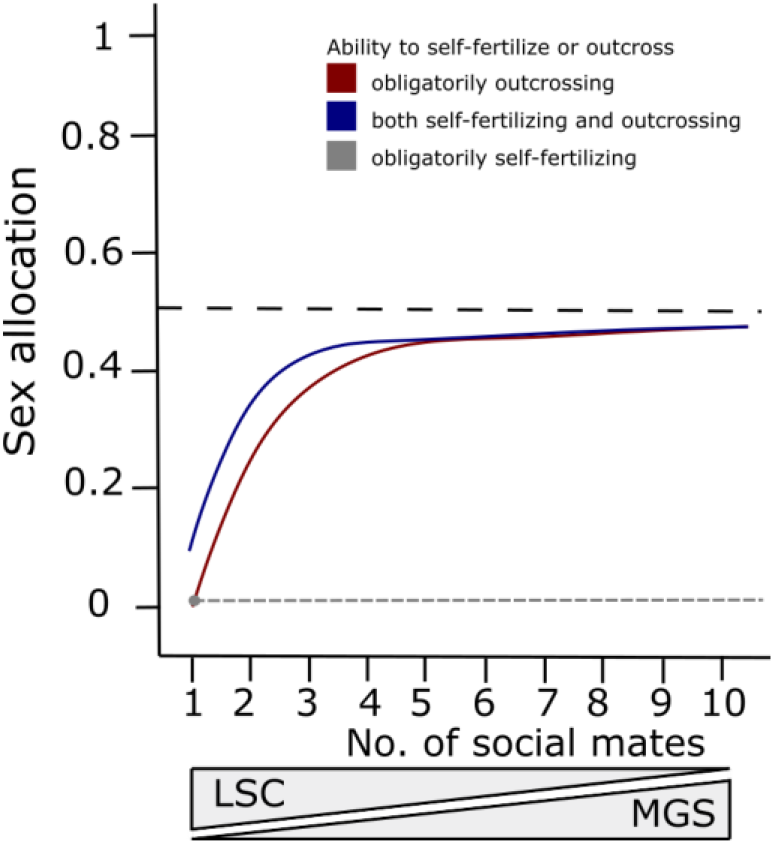
A visualization of our hypotheses for the effect of the number of social mates on the predicted sex allocation (i.e. the proportion of resources allocated to the male function) for species that either obligatorily outcross (red), species that both self-fertilize and outcross simultaneously (blue), or species that obligatorily self-fertilize (grey). As the number of social mates increases, the so-called mating group size (MGS, i.e. the number of actual mates plus one) is expected to increase, while the level of local sperm competition (LSC) is expected to decrease. Note that for species that obligatorily self-fertilize, mating group size always remains one and LSC is always maximal, leading to a highly female-biased sex allocation (i.e. the minimal male allocation to allow for full self-fertility), independently of the number of social mates (indicated by grey dot and stippled line). For species that both self-fertilize and outcross simultaneously, the MGS is already increased when the number of mates is one, since own sperm will compete with the partner’s sperm. And in species that outcross only, the prediction for when the number of mates is one is only the minimal male allocation to allow for full outcross-fertility. Note that these SA predictions are only approximate, since the degree to which MGS increases with the number of social mates will likely vary, and the extent of self-fertilization and outcrossing is unclear in the species that show both.

While most SA models investigate the effect of LSC on SA over evolutionary timescales, LSC can also vary temporally and spatially within an individual’s lifetime. In hermaphrodites, altering the SA in response to the current conditions can influence the immediate reproductive success of an individual, favouring the evolution of SA plasticity. Indeed, plastic SA has been suggested to be one of the advantages of hermaphroditism over separate-sex species (Charnov 1982; Michiels 1998, 1999). However, for species that do not often experience variation in LSC during their lifetime, we may not expect high levels of plasticity, particularly if there are costs to plasticity (DeWitt et al. 1998; Auld et al. 2010; Siljestam and Östman 2017).

Interestingly, while SA plasticity has now been documented in many hermaphroditic species (Schärer 2009), its evolution is still comparatively poorly understood, with few studies having investigated variation in SA plasticity across animal species in a controlled experimental context (Schleicherová et al. 2014). And finally, as SA can vary both between-species (e.g. due to differences in mating strategy over evolutionary timescales) and within-species (e.g. due to phenotypic plasticity), it is interesting to examine the relative magnitude of the interspecific versus intraspecific variation in SA.

An excellent model system for testing the effect of the mating strategy and the ability to self-fertilize on SA and SA plasticity is the free-living flatworm genus *Macrostomum* (Macrostomorpha, Platyhelminthes). This genus contains many species exhibiting one of at least two different mating strategies; one involving reciprocal mating and the other hypodermic insemination (Brand et al. 2021a; Schärer et al. 2011; Singh et al. in prep.). In reciprocally mating species, a facultative postcopulatory suck behaviour has been observed, in which an individual places its pharynx on top of its female antrum (the sperm-receiving and egg-laying organ) and appears to suck, and this has been hypothesised to remove received ejaculate components (Singh et al. in prep.; Schärer et al. 2004a; Vizoso et al. 2010). No such postcopulatory suck behaviour has been documented in hypodermically inseminating species (Singh et al. in prep.), which presumably exhibit forced unilateral mating, with sperm being hypodermically injected into the partner via a needle-like male copulatory organ (Schärer et al. 2011; Brand et al. 2021a). Moreover, both reciprocally mating and hypodermically inseminating species possess a female antrum, but in the latter this organ has a simple morphology and presumably serves only for egg-laying, while in the former it is usually more complex and used both for egg-laying and for receiving sperm from the partner (Schärer et al. 2011; Brand et al. 2021a). Interestingly, hypodermic insemination is also associated with a suite of morphological traits that potentially facilitate self-fertilization (Ramm et al. 2012, 2015; Giannakara and Ramm 2017), although self-fertilization has recently also been documented in at least one reciprocally mating species, *M. mirumnovem* (Singh et al. 2020b).

Here, we collected literature data on SA estimates from a range of experiments performed in six *Macrostomum* species, and we present additional data from a new and currently still undescribed species, *Macrostomum* sp. 22 (see Supplementary Information S1). In all studies the experimental design generally consisted in raising worms from juveniles in three different group sizes (isolated, pairs or octets) and then obtaining estimates of their SA once the worms had reached maturity. To facilitate SA plasticity comparisons between species, we calculated standardized effect sizes for SA plasticity in response to i) the presence of mating partners (i.e. comparing isolated worms vs. worms with partners) and ii) the strength of LSC (i.e. comparing paired worms vs. octet worms). Using SA estimates and these SA plasticity effect sizes, we then examined if the mating strategy and ability to self-fertilize predicted SA or SA plasticity, while accounting for the phylogenetic interrelationships. Lastly, considering the interspecific SA differences and the extensive SA plasticity shown by some of the species, we examined how much of the variation in SA occurs between-species versus within-species, by partitioning the SA variance into its interspecific and intraspecific components.

## Materials and Methods

### Study species

To examine how SA and SA plasticity evolves across the *Macrostomum* genus, we gathered data on SA and SA plasticity in seven *Macrostomum* species (we use the “Genus species Author, Year” format to refer to binomials and include citations to the relevant taxonomic works). Three species are hypodermically inseminating, namely *M. pusillum* Ax, 1951 (Ax 1951), *M. hystrix* Örsted, 1843 sensu Luther, 1905 (Örsted 1843; Luther 1905; Schärer et al. 2011), and the currently undescribed species *Macrostomum* sp. 22, with the SA data reported in, respectively, Giannakara and Ramm (2017), Winkler and Ramm (2018), and Supplementary Information S1. An additional four species are reciprocally mating, namely *M. janickei* Schärer, 2019, *M. cliftonense* Schärer and Brand, 2019, and *M. mirumnovem* Schärer and Brand, 2019 (Schärer et al. 2020; Zhang et al. 2021), with the SA data reported in Singh et al. (2020b), plus multiple studies in *M. lignano* Ladurner, Schärer, Salvenmoser, and Rieger, 2005 (Ladurner et al. 2005), with the SA data reported in several studies (see below). Details of all the SA studies are given in Supplementary Table S1.

The species *M. mirumnovem* (Singh et al. 2020b) and all three hypodermically inseminating species (Ramm et al. 2012, 2015; Giannakara and Ramm 2017, this study) can self-fertilize, and were classified accordingly. Moreover, we classified species as either reciprocally mating or hypodermically inseminating using both behavioural and morphological data. Behavioural data showed the presence of both reciprocal mating and the postcopulatory suck behaviour in *M. lignano, M. janickei, M. cliftonense*, and *M. mirumnovem*, while neither of these behaviours were observed in the hypodermically inseminating *M. pusillum, M. hystrix*, and *Macrostomum* sp. 22 (Singh et al., in prep.; Schärer et al. 2004a; Schärer et al. 2020). In addition, a classification of species using only morphological traits, which are known to be correlated with the mating strategy in *Macrostomum* (Singh et al. in prep.; Schärer et al. 2011), corroborated our above classification based on behaviour. Thus, species could also be classified morphologically as reciprocally mating or hypodermically inseminating depending on their male and female genital morphology, the sperm morphology, and the location of (received) allosperm (for details see Brand et al. 2021a).

### SA estimates and SA plasticity effect sizes across species

For most species the experimental procedure was similar to that described for *Macrostomum* sp. 22 (Supplementary Information S1), i.e., worms were raised from juveniles in three different group sizes (isolated, pairs or octets) until adulthood, when testis and ovary size was measured in one randomly chosen worm per replicate as a proxy for an individual’s male and female allocation. The SA was then estimated as testis size/(testis size+ovary size). In some experiments, the density of worms was varied independently of group size by using two enclosure sizes (small and large) (for details and sample sizes see Supplementary Table S1). We expect that the experiments are nevertheless comparable, since all experiments where density was included as a factor revealed that it did not have a significant effect on the estimate of SA.

For each experiment, we calculated the mean SA and respective standard deviation across the group sizes, weighted by the sample size for each group size, using the R package ‘Hmisc’ (Harrell 2020). To facilitate interspecies comparisons of the SA plasticity estimates, we calculated standardized effect sizes and their standard deviation, using Cohen’s d (Cohen 1988) with Hedges correction for small sample size (Hedges 1981), including confidence intervals (Howell 2011) of the effect sizes using the R package ‘effsize’ (Torchiano 2017). For each species, we calculated an effect size for SA plasticity to assess the effect of the presence of mating partners (i.e. comparing isolated worms vs. worms with partners) and the strength of LSC (i.e. comparing worms in pairs vs. worms in octets). Pairs represent a condition with high LSC whereas octets represent a condition with low LSC. Studies in *M. lignano* have shown that in larger groups individuals mate with many of the available partners (Janicke and Schärer 2009a; Janicke et al. 2013), that there is sperm displacement (Marie-Orleach et al. 2014) and that paternity is usually shared (Marie-Orleach et al. 2016; Vellnow et al. 2018).

For *M. lignano*, we collected datasets from multiple published experiments (up to 25^th^ November 2019), and calculated SA and SA plasticity effect sizes for each of them separately. In total, we found two and six studies, respectively, where we could extract the effects of the presence of mating partners (Schärer and Janicke 2009; Ramm et al. 2019) and the strength of LSC (Schärer and Ladurner 2003; Schärer et al. 2004b; Janicke and Schärer 2009b; Janicke et al. 2013; Marie-Orleach et al. 2014; Ramm et al. 2019). For Schärer and Ladurner (2003), we calculated SA from the available data, since that study only reported values for testis and ovary size. We excluded one previous study (Janicke and Schärer 2010) from our analysis, since the worms used in that study came from the same experiment as Janicke and Schärer (2009), and so these did not represent fully independent datasets (we used the latter study since it included more replicates). In addition, in Marie-Orleach et al. (2014) we combined the data from both inbred lines (HUB1 and DV1), since they did not differ in any of the traits measured. Also, note that in Schärer and Janicke (2009), there are only isolated and paired worms and hence the estimated effect size for the presence of mating partners does not include the effect of octets. This should not have a large effect on the calculated effect size, since in most cases the difference between the isolated and paired treatments is much larger than the difference between paired and octet treatments (Figure 2).

**Figure 2.**
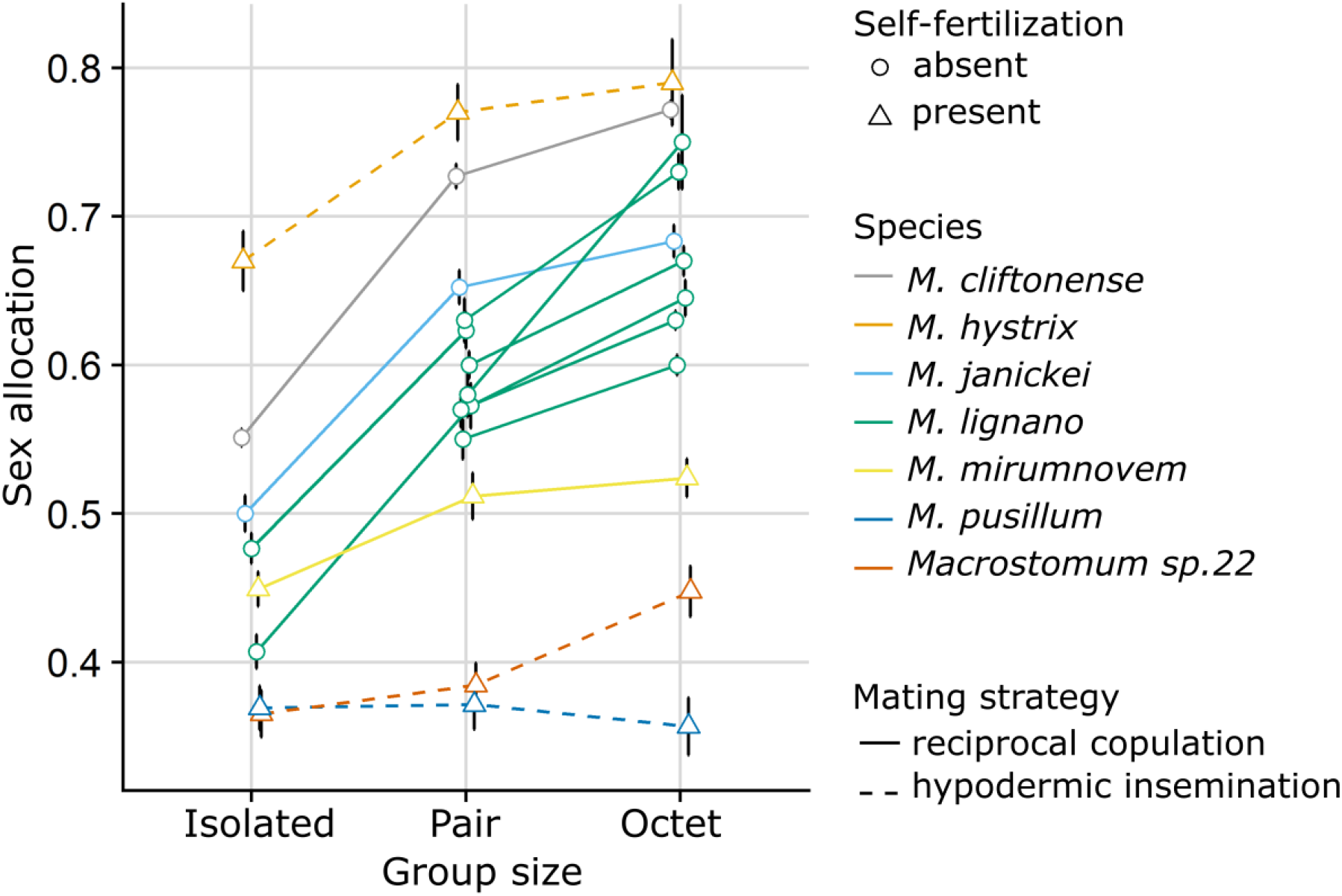
Effect of group size on estimates of sex allocation in seven different *Macrostomum* species (given by different colours). The line types represent the two mating strategies and the symbols represent the ability to self-fertilize. The plots show means and standard errors of the raw (untransformed) data. Note that for *M. lignano*, data from seven independent experiments are shown.

For *M. lignano*, a weighted mean across experiments was used to estimate the SA, and SA plasticity effect sizes (Turner and Bernard 2006), for both the presence of mating partners and the strength of LSC. For SA, we took the mean of the SA across experiments, weighted by the sample size of each experiment. To weight each effect size, it was multiplied by the inverse of its variance (inverse variance weight, IVW), which allowed us to take into consideration the differences in sample size across experiments, namely:

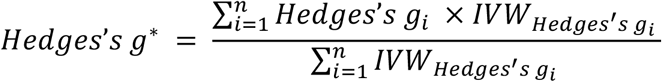

And its standard error was calculated as:

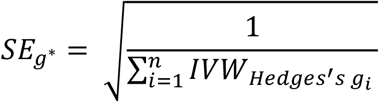

### Evolution of SA and SA plasticity

For the phylogenetically corrected analyses of the SA and SA plasticity effect sizes, we used a trimmed (ape package, Paradis et al. 2004) ultrametric phylogenomic tree constructed using an alignment containing 385 genes (and 74,175 variable sites) from 98 transcriptome-sequenced species, and calculated with a maximum likelihood approach (called H-IQ-TREE in Brand et al. 2021b). All bipartitions in this trimmed tree had maximal support (Figure 3), and trees constructed using a range of phylogenetic approaches had identical topologies with respect to the seven species studied here (Brand et al. 2021b), thus obviating the need to account for phylogenetic uncertainty in the following analyses.

**Figure 3.**
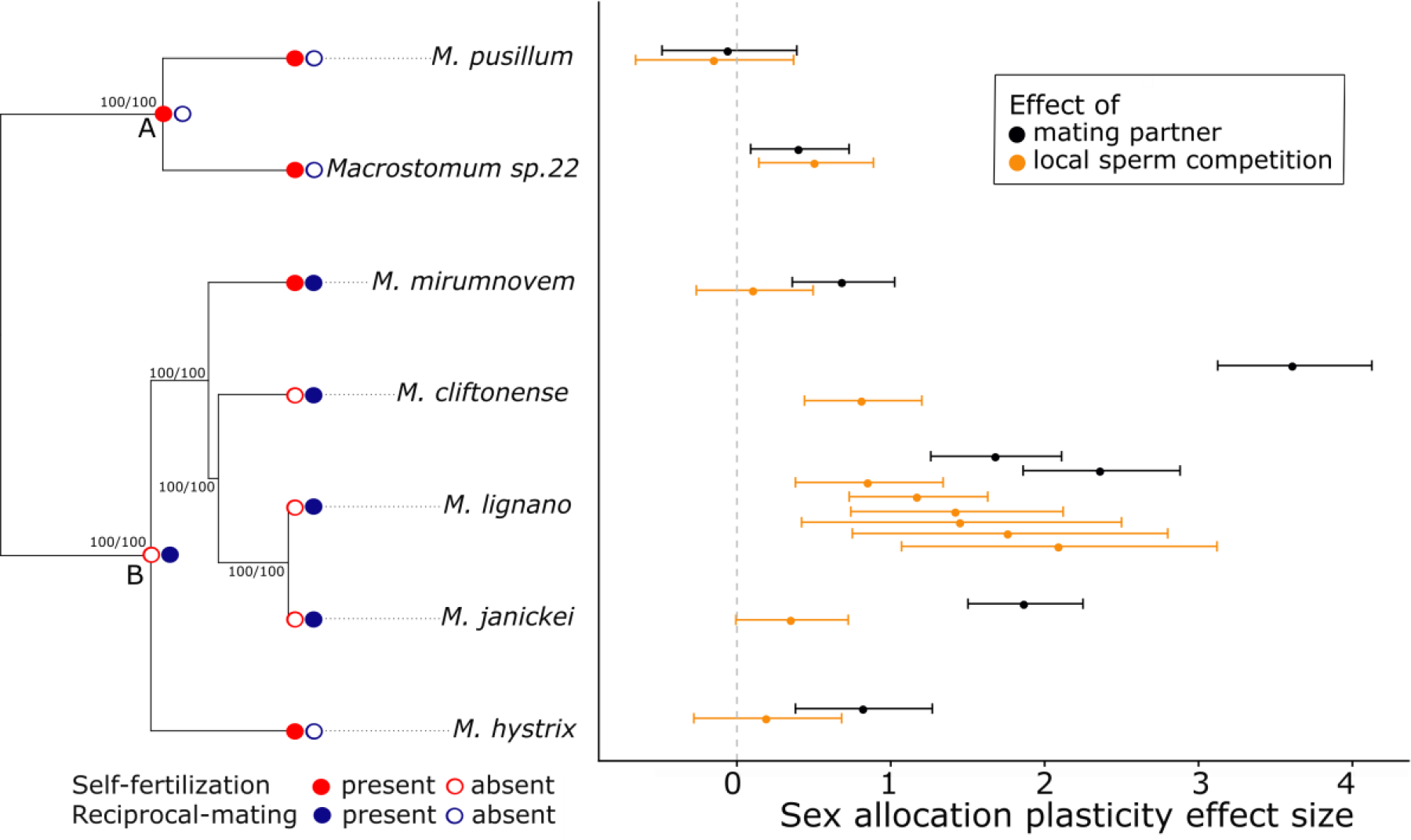
Standardized SA plasticity effect sizes for the effect of the presence of mating partners (i.e. comparing isolated worms vs. worms with partners) and the strength of local sperm competition (i.e. comparing worms in pairs vs. octets) among seven *Macrostomum* species (right side). The error bars represent 95% confidence intervals. For *M. lignano*, data from multiple experiments are shown. Also indicated is whether self-fertilization or/and reciprocal mating is present or absent in a species (left side), and these traits are mapped onto a trimmed maximum likelihood phylogeny of the genus (i.e. the H-IQ-TREE phylogeny from Brand et al. 2021b, which is based on 385 genes in 98 *Macrostomum* species), and all the shown bipartitions in this tree had maximal support, as indicated by ultrafast bootstrap support (first number) and approximate likelihood ratio tests (second number), respectively. A and B represent the inferred ancestral states at important internal nodes, suggesting that there are at least two independent origins of hypodermic insemination (Schärer et al. 2011; Brand et al. 2021a) and probably also self-fertilization in the genus (see also Methods and Supplementary Figure S1 for details).

Recent studies have shown that there are at least nine independent origins of hypodermic insemination in *Macrostomum* (Brand et al. 2021a). Thus *M. hystrix* clearly represents an independent origin of hypodermic insemination in an otherwise largely reciprocally mating clade, when compared to the other two hypodermically inseminating species, *M. pusillum* and *Macrostomum* sp. 22, which belong to a uniformly hypodermically mating clade (as indicated by the inferred ancestral states of the internal nodes in the phylogeny in Figure 3). Although a similarly unequivocal analysis for multiple independent origins of self-fertilization is currently not yet available, self-fertilization was not readily observed in multiple reciprocally mating species that we also cultured in the laboratory, including *M. spirale* Ax, 1956 (Ax 1956; Schärer et al. 2011), *M. axi* Papi, 1959 (Papi 1959), *M. clavituba* Ax, 2008 (Ax 2008), and *M. poznaniense* Kolasa, 1973 (Kolasa 1973), which were held in isolation for extended periods of time for behavoural experiments (P. Singh, personal observations). All of these species are members of the reciprocally mating clade and they fall outside of the subclade containing both *M. mirumnovem* and *M. hystrix* (Supplementary Figure S1). Thus, if we assume that the absence of self-fertilization is the most likely ancestral state in the entire reciprocally mating clade, then *M. hystrix* and *M. mirumnovem* represent at least one, or possibly two, independent origins of self-fertilization relative to that observed in *M. pusillum* and *Macrostomum* sp. 22 (Supplementary Figure S1).

As a preliminary analysis, we next examined if there was an association between self-fertilization and the mating strategy, since such an association would render these non-independent predictor variables. For this, we used the DISCRETE model in BayesTraits V.3.0.1 to test for correlated evolution between the presence of self-fertilization and the presence of hypodermic insemination using the Reversible Jump Markov Chain Monte Carlo (RJ MCMC) approach (Pagel 1994; Pagel and Meade 2006; Meade and Pagel 2016). To test support for correlated evolution, we compared the marginal likelihood of a dependent model, in which the presence of self-fertilization depends on the presence of hypodermic insemination, with an independent model, in which hypodermic insemination and self-fertilization evolve independently. We ran each RJ MCMC chain for ten million iterations, while discarding the first one million iterations as burn-in, after which the chain was sampled every 1000th iteration. We placed 1000 stepping stones (iterating each 10000 times), and used a gamma hyperprior (gamma 0 1 0 1) to obtain the marginal likelihood values for the models. We performed three separate runs for both the independent and dependent models to check for the stability of the likelihood values and convergence. We established that the chains had converged using Gelman and Rubin’s convergence diagnostic (Gelman and Rubin 1992) and that the effective sample size was >200 for all parameters, using the R package coda (Plummer et al. 2006). In addition, we also confirmed that the acceptance rate was usually between 20-40% (ideal when the chain is at convergence and indicating good mixing; Pagel and Meade 2006). We evaluated the alternative models using the Log Bayes Factor (BF) and used the convention that a BF value greater than 2 is considered as positive support for the best-fit model (Pagel and Meade 2006). We found only very weak support for the dependent model of evolution over the independent model for the association between self-fertilization and mating strategy, with all three runs for each model providing similar values (mean marginal likelihood, independent = -9.59, dependent = -9.57, BF: 0.05; see also Supplementary Table S2). This showed that the presence of self-fertilization appears uncorrelated to the mating strategy in our dataset, and we could therefore use the mating strategy and self-fertilization as independent predictors for our analysis of SA and SA plasticity.

We estimated the phylogenetic signal using Blomberg’s K (picante package version 1.8.2, Blomberg et al. 2003; Kembel et al. 2010) and Pagel’s λ (phytools package version 0.7-70, Pagel 1999; Revell 2012), for both SA and the two SA plasticity effect sizes. A phylogenetic signal value close to zero is suggestive of a trait evolving independently of the phylogeny, while a value close to one suggests that the traits evolve under Brownian motion. In our case, the estimates for SA showed a phylogenetic signal, while the two SA plasticity effect sizes did not differ significantly from 0 (Supplementary Table S3). Since our sample sizes are small, the likelihood ratio tests used to assess Pagel’s λ can be unreliable (Boettiger et al. 2012), as the asymptotic properties of maximum-likelihood estimation may not hold for small sample sizes. Thus, we present both phylogenetically corrected analyses, as well as analyses without correcting for phylogeny.

For the phylogenetically corrected analyses, we compared the fit of different character evolution models (i.e. Brownian motion, Ornstein–Uhlenbeck, and Early-burst) (geiger package, Harmon et al. 2008) for SA and the two SA plasticity effect sizes. While it has been suggested that datasets with small sample sizes can erroneously favour Ornstein–Uhlenbeck models over simpler models like Brownian motion (Cooper et al. 2016b,a), our small dataset was found to be more consistent with a Brownian motion model (see Supplementary Table S4). Thus, we used Brownian motion as the preferred model for the subsequent analysis. This does not necessarily imply, however, that the traits here actually evolve at random, but solely that a Brownian motion model fits our data better than the alternative Ornstein–Uhlenbeck or Early-burst models.

Using phylogenetic generalized least squares (PGLS) regressions and a Brownian motion model of character evolution (nlme package version 3.1-152, Pinheiro et al. 2014), we tested if the mating strategy (reciprocal mating vs. hypodermic insemination) or self-fertilization (presence or absence) predicted SA and the two SA plasticity effect sizes (presence of mating partners and strength of LSC). PGLS allows us to account for the phylogenetic non-independence of observations resulting from common evolutionary history of species. Furthermore, we also conducted an analysis without correcting for phylogeny, where we calculated Wilcoxon rank-sum tests to assess if the mating strategy or self-fertilization predicted the differences in SA or SA plasticity effect sizes. While comparative studies usually consider only the mean value of a trait, this does not allow us to incorporate intraspecific variability, which can be a possible source of error (Boucher et al. 2012; Tonnabel et al. 2018). Here, we therefore incorporated intraspecific variation for both the PGLS and Wilcoxon rank-sum tests using a resampling approach (with 10000 iterations). Each time we generated a random value from a normal distribution with the observed means and standard deviations for each species (Supplementary Table S1) and performed the analysis on this value. We report the mean values of PGLS in the main text and the distributions in Supplementary Figure S2.

Finally, to partition the observed variance in SA into its between-species and within-species components, we used the mean SA (or weighted mean SA across all experiments for *M. lignano*) for each group size and species (i.e. 3 observations per species), and fit a linear mixed model (lme4 package, Bates et al. 2015) using species as a random effect and calculated the percentage of variance explained by species. And additionally, to explore the impact that the variable outcomes of the different experiments in *M. lignano* might have on these estimates, we picked one mean SA value per group size and experiment at random, and then redid the above analysis (resampling 10000 times).

All statistical analyses were carried out using R version 4.0.5 (R Core Team), unless stated otherwise.

## Results

### SA and effect sizes of SA plasticity for all species

We found substantial variation in overall SA across the seven *Macrostomum* species (Figure 2, Supplementary Table S1), both with respect to the mating strategy (solid vs. stippled lines) and self-fertilization (circles vs. triangles). Two of the three hypodermically inseminating species (*Macrostomum* sp. 22 and *M. pusillum)*, exhibited a low SA. Similarly, and in line with our predictions, three of the four self-fertilizing species (*Macrostomum* sp. 22, *M. pusillum* and *M. mirumnovem*), exhibited a relatively low SA, likely indicating that SA in these species was female-biased. Interestingly, the exception to these patterns was *M. hystrix*, which had the highest SA of all the studied species. There was also interspecific variation in the effect size estimates of SA plasticity, even among relatively closely related species (Figure 3, Supplementary Table S1). For example, *M. cliftonense* and *M. lignano* exhibited the highest SA plasticity in response to the presence of mating partners and the strength of LSC, respectively. In addition, *M. lignano* exhibited SA plasticity across all experiments and while the magnitude varied somewhat, the confidence intervals overlapped, indicating that the estimates were fairly consistent across experiments (Figure 3).

### Evolution of SA and SA plasticity

The PGLS models showed that neither the mating strategy (mean: t_5_ = -0.56, P = 0.62) nor the ability to self-fertilize (mean: t_5_ = -0.54, P = 0.61) significantly predicted SA (Supplementary Figure S2a-d). And similar to SA, the mating strategy also did not significantly predict the SA plasticity effect size, neither due to the presence of mating partners (mean: t_5_ = 0.97, P = 0.38) nor the strength of LSC (mean: t_5_ = 0.29, P = 0.78) (Supplementary Figure S2e-f, S2i-j). In contrast, self-fertilization significantly predicted plasticity of SA in response to the presence of mating partners (mean: t_5_ = -2.98, P = 0.03, Supplementary Figure S2g-h), with this kind of plasticity being lower for selfing species (compare the black effect size estimates between selfing vs. non-selfing species in Figure 3). No significant association between self-fertilization and plasticity of SA was observed in response to the strength of LSC (mean: t_5_ = - 0.72, P = 0.52, Supplementary Figure S2k-l). Qualitatively similar results were obtained from the analyses that did not correct for phylogeny using Wilcoxon rank-sum tests (Supplementary Table S5).

Lastly, the linear-mixed model, using the weighed mean SA for *M. lignano*, showed that between-species variance explained 73.6% of the total SA variance (interspecific variance = 0.0177, and residual variance = 0.0063), suggesting that the observed SA variance between-species is nearly three times larger than the observed variance within-species (and the resampling approach showed very similar results, Supplementary Table S6).

## Discussion

Our study showed that there was substantial interspecific variation in both SA and SA plasticity in *Macrostomum*, also among closely related species, suggesting that both SA and SA plasticity are evolutionarily labile. Furthermore, while the mating strategy predicted neither SA nor the SA plasticity effect sizes, self-fertilizing species had a lower SA plasticity in response to the presence of mating partners. In the following, we discuss these findings in some detail.

In the context of phenotypic plasticity, very few studies have explored predictions of SA theory across multiple species in hermaphroditic animals (Hoch and Levinton 2012; Schleicherová et al. 2014), with most studies focussing solely on intraspecific comparisons (Schärer 2009). Hoch and Levinton (2012) tested Charnov’s (1980, 1982) mating group size model in two species of acorn barnacles, *Semibalanus balanoides* and *Balanus glandula*, by manipulating both the number and density of individuals in a natural setting. They showed that, while both species exhibit an increased male allocation (estimated using the sum of the mass of testes, sperm, seminal vesicle, and penis) at higher densities, there was no clear effect on SA from either treatment. In part this was because the species also responded in terms of their female allocation (estimated using egg mass), with both the number and density manipulation having a positive effect in *S. balanoides*—i.e. an effect in the opposite direction than predicted by the model—while there was no clear effect in *B. glandula*. The authors argued that this might result from interspecific differences in life-history traits. In contrast to the acorn barnacles, a study on resource allocation in response to different numbers of mating partners in three related polychaete worms, *Ophryotrocha diadema, O. adherens* and *O. gracilis*, showed that there was no significant plasticity in male allocation (estimated as the number of sperm produced by an individual) across the species (Schleicherová et al. 2014), while the species differed in the plasticity in female allocation (using multiple estimates, such as resource investment in eggs, total number of egg cocoons, and time interval between egg laying). The authors proposed that the magnitude of plasticity depended on the species-specific costs of the sex function, and also on the mating system of the species, and that there might be considerable resource allocation to behaviours linked to the male role in at least one of these species (Lorenzi et al. 2006; Santi et al. 2018). So, while both of these studies document interspecific variation in SA plasticity, they lacked the power to explore statistically whether interspecific differences in reproductive traits may affect the level of SA plasticity.

Our results showed that both SA and SA plasticity varied considerably across the studied *Macrostomum* species. Both hypodermically inseminating and self-fertilizing species tended to show a low (i.e. more female-biased) SA, with the notable exception of *M. hystrix*. This is interesting, since *M. hystrix* has been shown to represent an independent origin of hypodermic insemination in the reciprocally mating clade (Schärer et al. 2011). It has previously been suggested that SA could become more male-biased when a species shifts to hypodermic insemination, since hypodermic sperm might compete more in a fair-raffle-type sperm competition (Schärer and Janicke 2009), unless selfing were to become the dominant mating mode. Although capable of self-fertilization, *M. hystrix* is known to reach very high densities in the field (L. Schärer, pers. obs.), which could potentially favour the evolution of a high SA. In this context it is also important to note that our estimates of male and female allocation are not absolute, such that when we obtain SA estimates >0.5 that probably does not suggest a male-biased SA. This is because our SA estimate implicitly assumes that both testis and ovary area are similarly suitable and complete proxies for male and female reproductive allocation, respectively, across species. However, it is clear that these proxies do not provide absolute estimates of SA, for multiple reasons (see also Singh et al. 2020b). For example, the energetic expenditure per unit tissue could differ between the testes and ovaries (Schärer 2009), or there could be other components of male and female allocation that are not quantified by assessing gonad size, such as substantial provisioning of developing oocytes with yolk, sex-specific behaviours (Picchi and Lorenzi 2019), or the production of seminal fluid (Patlar et al. 2019), components that could themselves also be plastic.

We show that *M. lignano* and *M. janickei*, which are sibling species capable of hybridization (Singh et al. 2020a), exhibit differences in SA plasticity in response to the strength of LSC (although the 95% confidence intervals of two of the six *M. lignano* studies overlap somewhat with that of *M. janickei*). Variation in SA plasticity in species with similar reproductive biology could stem from different environmental conditions experienced by the species. For example, evolution of SA plasticity might not be favoured in species that inhabit relatively stable environments, especially if the maintenance of plasticity is costly (DeWitt et al. 1998; Auld et al. 2010; Siljestam and Östman 2017). Even in the absence of maintenance costs, there could still be production costs of plasticity, though if the benefits of adjusting SA outweigh these costs, then phenotypic plasticity could still be maintained. This cost-benefit ratio of SA plasticity can vary across species (Van Buskirk 2002; Steiner 2007) and environments (Ratikainen and Kokko 2019), leading to the retention or loss of plasticity in a given species. Moreover, such costs of SA plasticity might play an important role in both the evolution and the maintenance of simultaneous hermaphroditism (St. Mary 1997), and could also constrain plastic changes in SA, potentially leading to a suboptimal SA. A study in *O. diadema* showed that there were no large short-term fitness costs of sex adjustment for the species (Lorenzi et al. 2008), while in *M. lignano* production costs of SA plasticity have previously been documented (Sandner 2013), with an alternating group size environment leading to a lower hatchling production compared to a stable group size environment. We currently do not have data on production costs of plasticity in *M. janickei* when exposed to a stable vs. fluctuating environment.

Across all species, the estimates of plasticity effect sizes are larger for the presence of mating partners than for the strength of LSC (number of mating partners), and for *M. cliftonensis* and *M. janickei*, the confidence intervals of the two effect size estimates do not overlap. This suggests that the increase in SA going from paired to octet groups is not as drastic as going from isolated worms to worms in larger groups. A similar phenomenon has been observed in the freshwater snail, *Lymnaea stagnalis*, where paired snails showed higher expression of six seminal fluid protein (SFP) genes compared to isolated snails, while the SFP expression of snails in larger (6 individuals) groups was similar to that of paired snails (Nakadera et al. 2019). In our study, this situation could potentially arise if in the presence of multiple partners in octets, worms do not only increase their testis size, but also their rate of sperm production per unit testis size (Schärer and Vizoso 2007), which in *M. lignano* has been shown to occur due to an increase in the speed of spermatogenesis (Giannakara et al. 2016). If such effects occur generally (but see Giannakara & Ramm 2017 for *M. pusillum*, a species that lacks SA plasticity, see also below), we might underestimate the extent of plasticity in male allocation when solely measuring testis size. Alternatively, there could also be increased investment in other components of the male function (see above). A study in *O. diadema* showed that increased mating opportunities are accompanied by higher investment into behavioural components of male function, such as aggressive behaviour (Lorenzi et al. 2006; Santi et al. 2018).

The SA plasticity effect sizes did not differ significantly between species exhibiting the reciprocal and hypodermic mating strategies, while self-fertilizing species had a lower SA plasticity (in response to presence of mating partner, but not the strength of LSC). Interestingly, to date *M. mirumnovem* is the only reciprocally mating *Macrostomum* species in which self-fertilization has been recorded (Singh et al. 2020b), while studies have usually found self-fertilization in hypodermic species. Including *M. mirumnovem* in our study allowed us to disentangle the association between mating strategy and self-fertilization with respect to SA plasticity effect sizes, clearly demonstrating how including additional species in a phylogenetically-informed context leads to more informed interpretations.

Interesting questions that arise then concern the causes of the variation in SA and SA plasticity between the self-fertilizing species. An effect of self-fertilization on SA has been theoretically predicted, and while there has been both theoretical and empirical work exploring the effect of self-fertilization on SA in plants (Lemen 1980; Charlesworth and Charlesworth 1981; Schoen 1982; Charnov 1987; McKone 1987; Brunet 1992), there have been few such studies in animals (Johnston et al. 1998). A comparative study on SA in plants showed that resource allocation to flower parts can differ depending on the mating systems, such as the pollen-ovule ratio increasing with the likelihood of cross-pollination (Cruden 1977; but see Cruden and Lyon 1985).

In our study, one possible explanation for variation between self-fertilizing species could be if the species differ in their rate and pattern of self-fertilization, which could have an effect on both optimal SA and optimal SA plasticity. For example, in obligatorily self-fertilizing species, we would expect that they do not increase their SA with social group size, since LSC would be expected to remain high irrespective of the number of individuals in the local group (Figure 1). On the other hand, for species that exhibit facultative self-fertilization, we might expect considerable SA plasticity despite a low overall SA. Our results do conform somewhat to this pattern, with *M. pusillum*—hypothesised to be obligately self-fertilizing (Giannakara and Ramm 2017)—showing both the lowest overall SA and essentially no SA plasticity. Its closest congener, *Macrostomum* sp. 22, showed a similarly female-biased SA in isolated and paired worms, but significant SA plasticity between worms in pairs and octets. If worms shifted from self-fertilization when isolated, to exclusively outcrossing when in the presence of a mating partner, then the strength of LSC would not be expected to change from when they are alone to being in a pair. We would, however, expect the strength of LSC to drop from worms in pairs to worms in octets, favouring an increased in male allocation and SA, a scenario that would match the observed SA plasticity patterns in this species. This could suggest that *Macrostomum* sp. 22 may only self in isolation. In contrast, *M. hystrix—*thought to be a preferentially outcrossing species, with studies showing costs of self-fertilisation (Ramm et al. 2012, Giannakara and Ramm 2020) and delayed selfing in isolation, at least in some populations (Ramm et al. 2012)—was found to be more plastic than *M. pusillum*.

Finally, our results indicate that there appears to be considerably more variation in SA between-species than within-species in the sample of species we studied here, which is remarkable, since we saw extensive plasticity in SA in certain species in response to changes in group size, including across the multiple experiments in *M. lignano*. This result suggests that we can probably interpret variation in SA among field-collected worms of different *Macrostomum* species as being at least partially due to interspecific variation, even if they happen to have been sampled from different social environments, and may therefore also vary in part due to SA plasticity. Thus, although we cannot currently explain the observed interspecific variation in SA among the species studied here, our study will facilitate research in understanding the evolution of SA patterns across *Macrostomum* species, by allowing future studies to include SA estimates from field-collected worms.

Collectively, our results suggest that there is substantial interspecific variation in SA and SA plasticity in *Macrostomum*, with self-fertilization being a significant predictor for the latter (although the relatively small number of species we could include in this comparative study of course needs to be taken into consideration). Future studies should explore how the extent of self-fertilization, be it obligatorily or facultative, affects SA and SA plasticity using data from a greater number of species.

## Acknowledgements

We would like to thank Christian Felber for help with the morphological measurements of *Macrostomum* sp. 22, Lucas Marie-Orleach, Steven Ramm and Tim Janicke for providing access to their datasets, Dita Vizoso and Kaja Wasik for help with collecting specimens of *Macrostomum* sp. 22, Jeremias Brand, Axel Wiberg and Nikolas Vellnow for helpful discussions, and Gudrun Viktorin, Daniel Lüscher, Jürgen Hottinger, Lukas Zimmerman and Urs Stiefel for technical support. This research was supported by grants 31003A_162543 and 310030_184916 of the Swiss National Science Foundation (SNSF) to LS.

## References

Auld, J. R., A. A. Agrawal, and R. A. Relyea. 2010. Re-evaluating the costs and limits of adaptive phenotypic plasticity. Proceedings of the Royal Society B: Biological Sciences 277:503–511.

Ax, P. 1951. Die Turbellarien des Eulitorals der Kieler Bucht. Zoologische Jahrbuecher Abteilung fuer Systematik Oekologie und Geographie der Tiere 80:277–378.

Ax, P. 1956. Les Turbellariés des étangs côtiers du littoral méditerranéen de la France méridionale… Hermann et Cie.

Ax, P. 2008. Plathelminthes aus Brackgewässern der Nordhalbkugel. Abhandlungen der Mathematisch-naturwissenschaftligen Klasse 2008:1.

Bates, D., M. Mächler, B. M. Bolker, and S. C. Walker. 2015. Fitting linear mixed-effects models using lme4. Journal of Statistical Software, doi: 10.18637/jss.v067.i01.

Blomberg, S. P., T. Garland, and A. R. Ives. 2003. Testing for phylogenetic signal in comparative data: behavioral traits are more labile. Evolution 57:717–745.

Boettiger, C., G. Coop, and P. Ralph. 2012. Is your phylogeny informative? Measuring the power of comparative methods. Evolution 66:2240–2251.

Boucher, F. C., W. Thuiller, C. Roquet, R. Douzet, S. Aubert, N. Alvarez, and S. Lavergne. 2012. Reconstructing the origins of high-alpine niches and cushion life form in the genus Androsace S.L. (Primulaceae). Evolution 66:1255–1268.

Brand, J. N., L. J. Harmon, and L. Schärer. 2021a. Frequent origins of traumatic insemination involve convergent shifts in sperm and genital morphology. BioRxiv doi: 10.1101/2021.02.16.431427.

Brand, J. N., G. Viktorin, R. A. W. Wiberg, C. Beisel, and L. Schärer. 2021b. Large-scale phylogenomics of the genus Macrostomum (Platyhelminthes) reveals cryptic diversity and novel sexual traits. In press. Molecular Phylogenetics and Evolution. BioRxiv doi: 2021.03.28.437366.

Brunet, J. 1992. Sex allocation in hermaphroditic plants. Trends in Ecology & Evolution 7:79–84.

Campbell, D. R. 2000. Experimental tests of sex-allocation theory in plants. Trends in Ecology & Evolution 15:227–232.

Charlesworth, D., and B. Charlesworth. 1981. Allocation of resources to male and female functions in hermaphrodites. Biological Journal of the Linnean Society 15:57–74.

Charnov, E. L. 1987. On sex allocation and selfing in higher plants. Evolutionary Ecology 1:30–36.

Charnov, E. L. 1980. Sex allocation and local mate competition in barnacles. Marine Biology Letters 1:269–272.

Charnov, E. L. 1979. Simultaneous hermaphroditism and sexual selection. Proceedings of the National Academy of Sciences 76:2480–2484.

Charnov, E. L. 1996. Sperm competition and sex allocation in simultaneous hermaphrodites. Evolutionary Ecology 10:457–462.

Charnov, E. L. 1982. The theory of sex allocation. Monographs in population biology 18:1–355. Princeton University Press, Cambridge.

Cohen, J. 1988. Statistical power analysis for the behavioral sciences.

Cooper, N., G. H. Thomas, and R. G. FitzJohn. 2016a. Shedding light on the ‘dark side’ of phylogenetic comparative methods. Methods in Ecology and Evolution 7:693–699.

Cooper, N., G. H. Thomas, C. Venditti, A. Meade, and R. P. Freckleton. 2016b. A cautionary note on the use of Ornstein Uhlenbeck models in macroevolutionary studies. Biological Journal of the Linnean Society 118:64–77.

Cruden, R. W. 1977. Pollen-Ovule Ratios: A Conservative Indicator of Breeding Systems in Flowering Plants. Evolution 31.

Cruden, R. W., and D. L. Lyon. 1985. Patterns of biomass allocation to male and female functions in plants with different mating systems. Oecologia 66.

DeWitt, T. J., A. Sih, and D. S. Wilson. 1998. Costs and limits of phenotypic plasticity. Trends in Ecology & Evolution 13:77–81.

Friedman, J., and S. C. H. Barrett. 2011. Genetic and environmental control of temporal and size-dependent sex allocation in a wind-pollinated plant. Evolution 65:2061–2074.

Gelman, A., and D. B. Rubin. 1992. Inference from Iterative Simulation Using Multiple Sequences. Statistical Science 7:457–472.

Giannakara, A., and S. A. Ramm. 2020. Evidence for inter-population variation in waiting times in a self-fertilizing flatworm. Invertebrate Reproduction & Development 64:158–168.

Giannakara, A., and S. A. Ramm. 2017. Self-fertilization, sex allocation and spermatogenesis kinetics in the hypodermically inseminating flatworm Macrostomum pusillum. The Journal of Experimental Biology 220:1568–1577.

Giannakara, A., L. Schärer, and S. A. Ramm. 2016. Sperm competition-induced plasticity in the speed of spermatogenesis. BMC Evolutionary Biology 16:60.

Greeff, J. M., and N. K. Michiels. 1999. Sperm Digestion and Reciprocal Sperm Transfer Can Drive Hermaphrodite Sex Allocation to Equality. The American Naturalist 153:421–430.

Guo, Q., D. G. Brockway, and X. Chen. 2017. Temperature-related sex allocation shifts in a recovering keystone species, Pinus palustris. Plant Ecology & Diversity 10:303–310.

Hamilton, W. D. 1967. Extraordinary sex ratios. Science 156:477–488.

Harmon, L. J., J. T. Weir, C. D. Brock, R. E. Glor, and W. Challenger. 2008. GEIGER: investigating evolutionary radiations. Bioinformatics 24:129–131.

Harrell, F. E. 2020. Hmisc: Harrell miscellaneous. R package version 4.4-0.

Hedges, L. V. 1981. Distribution theory for glass’s estimator of effect size and related estimators. Journal of Educational Statistics 6:107–128.

Hoch, J. M., and J. S. Levinton. 2012. Experimental tests of sex allocation theory with two species of simultaneously hermaphroditic acorn barnacles. Evolution 66:1332–1343.

Howell, D. C. 2011. Confidence intervals on effect size. Univresity of Vermont, doi: 10.1134/S0362119713030110.

Janicke, T., L. Marie-Orleach, K. De Mulder, E. Berezikov, P. Ladurner, D. B. Vizoso, and L. Schärer. 2013. Sex allocation adjustment to mating group size in a simultaneous hermaphrodite. Evolution 67:3233–3242.

Janicke, T., and L. Schärer. 2009a. Determinants of mating and sperm-transfer success in a simultaneous hermaphrodite. Journal of Evolutionary Biology 22:405–415.

Janicke, T., and L. Schärer. 2009b. Sex allocation predicts mating rate in a simultaneous hermaphrodite. Proceedings of the Royal Society B: Biological Sciences 276:4247–4253.

Janicke, T., and L. Schärer. 2010. Sperm competition affects sex allocation but not sperm morphology in a flatworm. Behavioral Ecology and Sociobiology 64:1367–1375.

Johnston, M. O., B. Das, and W. R. Hoeh. 1998. Negative correlation between male allocation and rate of self-fertilization in a hermaphroditic animal. Proceedings of the National Academy of Sciences 95:617–620.

Kembel, S. W., P. D. Cowan, M. R. Helmus, W. K. Cornwell, H. Morlon, D. D. Ackerly, S. P. Blomberg, and C. O. Webb. 2010. Picante: R tools for integrating phylogenies and ecology. Bioinformatics 26:1463–1464.

Kolasa, J. 1973. Two New Species of Macrostomum (Turbellaria), A Redescription of an Established Species and New Records from Poland. Bolletino di zoologia 40:181–200.

Ladurner, P., L. Schärer, W. Salvenmoser, and R. M. Rieger. 2005. A new model organism among the lower Bilateria and the use of digital microscopy in taxonomy of meiobenthic Platyhelminthes: Macrostomum lignano, n. sp. (Rhabditophora, Macrostomorpha). Journal of Zoological Systematics and Evolutionary Research 43:114–126.

Lange, R., K. Reinhardt, N. K. Michiels, and N. Anthes. 2013. Functions, diversity, and evolution of traumatic mating. Biological Reviews 88:585–601.

Lemen, C. 1980. Allocation of reproductive effort to the male and female strategies in wind-pollinated plants. Oecologia 45:156–159.

Lloyd, D. G. 1984. Gender allocaations in outcrossing cosexual plants. Pp. 277–300 in R. Dirzo and J. Sarukhan, eds. Perspectives in Plant Population Ecology. Sinauer, Sunderland, MA.

Lopez, S., and C. A. Dominguez. 2003. Sex choice in plants: facultative adjustment of the sex ratio in the perennial herb Begonia gracilis. Journal of Evolutionary Biology 16:1177–1185.

Lorenzi, M. C., D. Schleicherová, and G. Sella. 2006. Life history and sex allocation in the simultaneously hermaphroditic polychaete worm Ophryotrocha diadema: The role of sperm competition. Integrative and Comparative Biology 46:381–389.

Lorenzi, M. C., D. Schleicherova, and G. Sella. 2008. Sex adjustments are not functionally costly in simultaneous hermaphrodites. Marine Biology 153:599–604.

Luther, A. 1905. Zur Kenntnis der Gattung Macrostoma. Festschrift für Palmén, Helsingfors 5:1-61, 64 plates.

Marie-Orleach, L., T. Janicke, D. B. Vizoso, P. David, and L. Schärer. 2016. Quantifying episodes of sexual selection: Insights from a transparent worm with fluorescent sperm. Evolution 70:314–328.

Marie-Orleach, L., T. Janicke, D. B. Vizoso, M. Eichmann, and L. Schärer. 2014. Fluorescent sperm in a transparent worm: validation of a GFP marker to study sexual selection. BMC evolutionary biology 14:148.

McKone, M. J. 1987. Sex Allocation and Outcrossing Rate: A Test of Theoretical Predictions Using Bromegrasses (Bromus). Evolution 41:591.

Meade, A., and M. Pagel. 2016. Bayes Traits.

Michiels, N. K. 1998. Mating Conflicts and Sperm Competition in Simultaneous Hermaphrodites. Pp. 219–254 in Sperm Competition and Sexual Selection. Elsevier.

Michiels, N. K. 1999. Sexual adaptations to high density in hermaphrodites. Invertebrate Reproduction & Development 36:35–40.

Nakadera, Y., A. Giannakara, and S. A. Ramm. 2019. Plastic expression of seminal fluid protein genes in a simultaneously hermaphroditic snail. Behavioral Ecology 30:904–913.

Örsted, A. S. 1843. Forsog til en ny classification af Planarierne (Planariea Duges) grundet paa mikroskopisk-anatomiske Undersogelser. Naturhistorisk Tidsskrift 4:519–581.

Pagel, M. 1994. Detecting correlated evolution on phylogenies: a general method for the comparative analysis of discrete characters. Proceedings of the Royal Society of London. Series B: Biological Sciences 255:37–45.

Pagel, M. 1999. Inferring the historical patterns of biological evolution. Nature 401:877–884.

Pagel, M., and A. Meade. 2006. Bayesian Analysis of Correlated Evolution of Discrete Characters by Reversible-Jump Markov Chain Monte Carlo. The American Naturalist 167:808–825.

Papi, F. 1959. Specie nuove o poco note del gen. Macrostomum (Turbellaria: Macrostomida) rinvenute in Italia. Monitore Zoologico Italiano 66:84–102.

Paradis, E., J. Claude, and K. Strimmer. 2004. APE: Analyses of Phylogenetics and Evolution in R language. Bioinformatics 20:289–290.

Parker, G. A. 1970. Sperm competition and its evolutionary consequences in the insects. Biological Reviews 45:525–567.

Parker, G. A. 1998. Sperm Competition and the Evolution of Ejaculates: Towards a Theory Base. Pp. 3–54 in Sperm competition and Sexual selection.

Patlar, B., M. Weber, and S. A. Ramm. 2019. Genetic and environmental variation in transcriptional expression of seminal fluid proteins. Heredity 122:595–611.

Pen, I., and F. J. Weissing. 1999. Sperm competition and sex allocation in simultaneous hermaphrodites: A new look at Charnov’s invariance principle. Evolutionary Ecology Research 517–525.

Picchi, L., and M. C. Lorenzi. 2019. Gender-related behaviors: evidence for a trade-off between sexual functions in a hermaphrodite. Behavioral Ecology 30:770–784.

Pinheiro, J., D. Bates, S. DebRoy, and D. Sarkar. 2014. R Core Team (2014). nlme: linear and nonlinear mixed effects models. R package version 3.1–117. URL: http://cran.r-project.org/web/packages/nlme/index.html.

Plummer, M., N. Best, K. Cowles, and K. Vines. 2006. {CODA}: Convergence Diagnosis and Output Analysis for {MCMC}. R News.

Rademaker, M. C. J., and T. J. Jong. 1999. The Shape of the Female Fitness Curve for Cynoglossum officinale: Quantifying Seed Dispersal and Seedling Survival in the Field. Plant Biology 1:351–356.

Ramm, S. A., B. Lengerer, R. Arbore, R. Pjeta, J. Wunderer, A. Giannakara, E. Berezikov, P. Ladurner, and L. Schärer. 2019. Sex allocation plasticity on a transcriptome scale: Socially sensitive gene expression in a simultaneous hermaphrodite. Molecular Ecology 28:2321–2341.

Ramm, S. a, A. Schlatter, M. Poirier, and L. Schärer. 2015. Hypodermic self-insemination as a reproductive assurance strategy. Proceedings of the Royal Society B: Biological Sciences 282:20150660.

Ramm, S. A., D. B. Vizoso, and L. Schärer. 2012. Occurrence, costs and heritability of delayed selfing in a free-living flatworm. Journal of Evolutionary Biology 25:2559–2568.

Ratikainen, I. I., and H. Kokko. 2019. The coevolution of lifespan and reversible plasticity. Nature Communications 10:538.

Reinhardt, K., N. Anthes, and R. Lange. 2015. Copulatory wounding and traumatic insemination. Cold Spring Harbor Perspectives in Biology 7:a017582.

Revell, L. J. 2012. phytools: an R package for phylogenetic comparative biology (and other things). Methods in Ecology and Evolution 3:217–223.

Rosas, F., and C. A. Domínguez. 2009. Male sterility, fitness gain curves and the evolution of gender specialization from distyly in Erythroxylum havanense. Journal of Evolutionary Biology 22:50–59.

Sandner, P. 2013. Impacts of sperm competition on mating behaviour and life history traits in a simultaneous hermaphrodite. University of Basel.

Santi, M., L. Picchi, and M. C. Lorenzi. 2018. Dynamic modulation of reproductive strategies in a simultaneous hermaphrodite and preference for the male role. Animal Behaviour 146:87–96.

Schärer, L. 2009. Tests of sex allocation theory in simultaneously hermaphroditic animals. Evolution 63:1377–1405.

Schärer, L., J. N. Brand, P. Singh, K. S. Zadesenets, C.-P. Stelzer, and G. Viktorin. 2020. A phylogenetically informed search for an alternative Macrostomum model species, with notes on taxonomy, mating behavior, karyology, and genome size. Journal of Zoological Systematics and Evolutionary Research 58:41–65.

Schärer, L., and T. Janicke. 2009. Sex allocation and sexual conflict in simultaneously hermaphroditic animals. Biology Letters 5:705–708.

Schärer, L., T. Janicke, and S. A. Ramm. 2015. Sexual conflict in hermaphrodites. Cold Spring Harbor Perspectives in Biology 7:a017673.

Schärer, L., G. Joss, and P. Sandner. 2004a. Mating behaviour of the marine turbellarian Macrostomum sp.: these worms suck. Marine Biology 145:373–380.

Schärer, L., and P. Ladurner. 2003. Phenotypically plastic adjustment of sex allocation in a simultaneous hermaphrodite. Proceedings of the Royal Society B: Biological Sciences 270:935–941.

Schärer, L., P. Ladurner, and R. M. Rieger. 2004b. Bigger testes do work more: experimental evidence that testis size reflects testicular cell proliferation activity in the marine invertebrate, the free-living flatworm Macrostomum sp. Behavioral Ecology and Sociobiology 56:420–425.

Schärer, L., D. T. J. Littlewood, A. Waeschenbach, W. Yoshida, and D. B. Vizoso. 2011. Mating behavior and the evolution of sperm design. Proceedings of the National Academy of Sciences 108:1490–1495.

Schärer, L., and I. Pen. 2013. Sex allocation and investment into pre-and post-copulatory traits in simultaneous hermaphrodites: the role of polyandry and local sperm competition. Philosophical transactions of the Royal Society of London. Series B, Biological sciences 368:20120052.

Schärer, L., and D. B. Vizoso. 2007. Phenotypic plasticity in sperm production rate: There’s more to it than testis size. Evolutionary Ecology 21:295–306.

Schleicherová, D., G. Sella, S. Meconcelli, R. Simonini, M. P. Martino, P. Cervella, and M. C. Lorenzi. 2014. Does the cost of a function affect its degree of plasticity? A test on plastic sex allocation in three closely related species of hermaphrodites. Journal of Experimental Marine Biology and Ecology 453:148–153.

Schoen, D. J. 1982. Male reproductive effort and breeding system in an hermaphroditic plant. Oecologia 53:255–257.

Siljestam, M., and Ö. Östman. 2017. The combined effects of temporal autocorrelation and the costs of plasticity on the evolution of plasticity. Journal of Evolutionary Biology 30:1361–1371.

Singh, P., D. N. Ballmer, M. Laubscher, and L. Schärer. 2020a. Successful mating and hybridisation in two closely related flatworm species despite significant differences in reproductive morphology and behaviour. Scientific Reports 10:12830.

Singh, P., J. N. Brand, and L. Schärer. n.d. Evolution of the suck behaviour, a postcopulatory female resistance trait in a hermaphroditic flatworm genus.

Singh, P., N. Vellnow, and L. Schärer. 2020b. Variation in sex allocation plasticity in three closely related flatworm species. Ecology and Evolution 10:26–37.

St. Mary, C. M. 1997. Sequential Patterns of Sex Allocation in Simultaneous Hermaphrodites: Do We Need Models That Specifically Incorporate This Complexity? The American Naturalist 150:73–97.

Steiner, U. K. 2007. Investment in defense and cost of predator-induced defense along a resource gradient. Oecologia 152:201–210.

Tonnabel, J., F. M. Schurr, F. Boucher, W. Thuiller, J. Renaud, E. J. P. Douzery, and O. Ronce. 2018. Life-History Traits Evolved Jointly with Climatic Niche and Disturbance Regime in the Genus Leucadendron (Proteaceae). The American Naturalist 191:220–234.

Torchiano, M. 2017. Package “effsize”: efficient effect size computation. CRAN Package, doi: 10.5281/ZENODO.196082.

Turner, H. M., and R. M. Bernard. 2006. Calculating and Synthesizing Effect Sizes. Contemporary Issues in Communication Science and Disorders 33:42–55.

Van Buskirk, J. 2002. A Comparative Test of the Adaptive Plasticity Hypothesis: Relationships between Habitat and Phenotype in Anuran Larvae. The American Naturalist 160:87–102.

van Velzen, E., L. Schärer, and I. Pen. 2009. The effect of cryptic female choice on sex allocation in simultaneous hermaphrodites. Proceedings of the Royal Society B: Biological Sciences 276:3123–3131.

Vellnow, N., L. Marie-Orleach, K. S. Zadesenets, and L. Schärer. 2018. Bigger testes increase paternity in a simultaneous hermaphrodite, independently of the sperm competition level. Journal of Evolutionary Biology 31:180–196.

Vizoso, D. B., G. Rieger, and L. Schärer. 2010. Goings-on inside a worm: Functional hypotheses derived from sexual conflict thinking. Biological Journal of the Linnean Society 99:370–383.

Winkler, L., and S. A. Ramm. 2018. Experimental evidence for reduced male allocation under selfing in a simultaneously hermaphroditic animal. Biology Letters 14:20180570.

Zhang, S., Y. Shi, Z. Zeng, F. Xin, L. Deng, and A. Wang. 2021. Two New Brackish-Water Species of Macrostomum (Platyhelminthes: Macrostomorpha) from China and Their Phylogenetic Positions. Zoological Science 38.

